# Patient-Centric Development of a Low-Volume, Low-Viscosity, Once Every 4 Weeks (Q4W) Autoinjector Combination Product for Povetacicept

**DOI:** 10.64898/2026.10.20.760382

**Authors:** Joanie Lucas, J. Alaina Floyd, Erin Hickey, Christopher Perez-Lopes, Ryan Lehane, Meera Chawla, Mike Salema, Gabriel Graffius

## Abstract

Biologic therapies delivered subcutaneously (SC) are increasingly moving toward at-home self-administration using drug-device combination products. This shift places greater emphasis on the patient experience, and factors such as dose volume, injection time, dosing frequency, viscosity, injection force, and ease of use can affect treatment burden. This article focuses on the impact of balancing high protein concentration with low viscosity during drug product formulation development for an improved patient experience for povetacicept, an investigational, subcutaneously administered dual B-cell activating factor (BAFF) and a proliferation-inducing ligand (APRIL) inhibitor being developed for the treatment of multiple kidney diseases, including IgA nephropathy (IgAN). For povetacicept, formulation and device considerations were integrated throughout development. Studies evaluated protein concentration, dose volume, viscosity, injection force, injection time, and usability, in addition to standard stability, to develop a low-volume Q4W autoinjector. The selected 175 mg/mL formulation delivers an 80 mg dose in 0.46 mL. The viscosity was no more than 10 cP at 15–25°C and in a 1 mL prefilled syringe, the average glide force was 6 N, and the injection time at a constant force of 20 N was < 2 seconds. The same prefilled syringe was incorporated into an autoinjector with an injection time of less than 2 seconds in air. We conducted a limited assessment of an approved treatment for IgAN (Voyxact) administered via prefilled syringe (PFS) for effects of viscosity, fill volume and syringe configuration under the same experimental conditions. The viscosity at room temperature was 16 cP, with an average glide force of 27N and injection time of 19 seconds for the 2 mL dose.

In conclusion, povetacicept was designed to support convenient and user-friendly at-home self-administration through Q4W dosing, a low injection volume, a low viscosity formulation, rapid delivery, and an easy-to-use autoinjector. This integrated formulation and device development approach demonstrates how patient-centric considerations can be incorporated in combination product development. Together, these attributes have the potential to reduce the practical burden associated with repeated, long-term treatment for IgAN patients.

## Introduction

For chronic diseases requiring long-term treatment, drug administration characteristics can meaningfully influence the patient’s experience. As biologic therapies increasingly transition to self-administration, several factors including dose volume, injection time, dosing frequency, viscosity, injection force, and general device usability have become important considerations in product development.^1,2^

Subcutaneous administration offers advantages for patients by enabling treatment outside of healthcare settings;^3^ however, development of subcutaneous biologics can present formulation challenges. Therapeutic doses often necessitate highly concentrated protein formulations to maintain practical injection volumes for SC administration, but increasing protein concentration can increase solution viscosity and product instability.^4,5^ In turn, elevated viscosity may increase the force required for administration, prolong injection times, and restrict delivery-device options.^6^ Consequently, formulation development must balance protein concentration, stability, and injectability in order to achieve a patient-friendly administration experience. This can increase the confidence and willingness of patients to self-administer, thereby reducing treatment burden and increasing patient adherence.

The close relationship between formulation characteristics and device performance requires the development of the drug product and the delivery system in parallel. As self-administered therapies become more common,^7^ there has been an increasing emphasis on the development of delivery devices that minimize administration burden and simplify the injection process. Autoinjectors have emerged as an attractive option because they automate key steps of dose administration and can be easier to use for patients with dexterity limitations and are commonly used in the delivery of peptides and biologics for rheumatology, dermatology, and metabolic (e.g., GLP-1 based therapies) indications.^8,9^ The use of an autoinjector, however, depends in part on formulation characteristics that enable rapid and reliable dose delivery.^10^

IgA nephropathy (IgAN) is a serious, chronic, immune-mediated kidney disease for which newer therapies are emerging. A recent qualitative survey study of adult patients with IgAN in the US revealed that 96% of those interviewed (N=24 of 25) preferred an autoinjector relative to injection via a prefilled syringe (PFS), and 80% (N=20 of 25) preferred Q4W versus Q1W dosing.^11^ Povetacicept is an investigational, subcutaneously administered dual B-cell activating factor (BAFF) and A proliferation-inducing ligand (APRIL) inhibitor being developed for the treatment of multiple diseases, including IgA nephropathy (IgAN).^12^ Development of the povetacicept drug-device combination product was guided by a patient-centered strategy aimed at optimizing both the formulation and delivery system to support convenient Q4W subcutaneous administration using an autoinjector. This article describes the scientific and engineering considerations underlying the development of the povetacicept drug-autoinjector combination, including formulation optimization, injectability assessments, and administration-related attributes relevant to the patient experience. In addition, a limited assessment of an approved commercial IgAN product (Voyxact) administered via PFS was included to provide descriptive external context for the effects of viscosity, fill volume, and syringe configuration under the experimental conditions used.

## Materials

Polyethylene glycol (PEG 400 molecular weight) and purified water were combined to prepare nominal viscosity standards of 5, 10, 15, and 20 cP. The verified viscosities were 5.4, 10.0, 16.3, and 21.9 cP, respectively.

Injectability testing was performed using Becton Dickinson Neopak 1 mL long and 2.25 mL borosilicate glass prefilled syringes (PFS). Both configurations used a 27-gauge, ½-inch, 5-bevel stainless-steel needle and a FluoroTec-coated bromobutyl rubber stopper. Syringe internal diameters were 6.35 mm for the 1 mL configuration and 8.65 mm for the 2.25 mL configuration.

Povetacicept viscosity data was obtained from formulation screening studies. Combination product injectability studies were conducted with 1 mL PFS (80 mg/0.46 mL) and 2.25 mL PFS (240 mg/1.37 mL) obtained from long-term storage inventory. Povetacicept autoinjector performance data were obtained from development and clinical batch release and stability data. Commercially-available Voyxact (sibeprenlimab-szsi) prefilled syringes were obtained through an authorized source and were used for a limited laboratory assessment.

## Methods

### Drug Product Viscosity Measurements

During formulation development, povetacicept samples were analyzed with an Anton Paar MCR92 cone-and-plate rheometer with a CP20-0.5 cone. A 25 µL sample was measured over shear rates from 50 to 2000 s^−1^; viscosity values were averaged over the shear-rate range of 500–1000 s^−1^. Triplicate measurements were performed using 25 µL per measurement, unless otherwise noted.

### PEG Samples Viscosity Verification

Dynamic viscosity of PEG viscosity standards was measured on a DV Next LV cone-and-plate rotational viscometer (Brookfield/AMETEK) fitted with a CPA-51Z spindle, with the cone–plate gap set to 0.0005. Sample temperature was controlled with a TC-650 circulating water bath set to 25 °C. Measurements were performed using 0.5 mL samples at a rotational speed of 60 rpm with a run time of 30 s. Prior to each test session, system performance was verified using 5 cP and 20 cP mineral oil viscosity standards.

### Break Loose and Glide Force Testing

Break Loose and Glide Force (BLGF) testing was performed using an Instron 68SC-1 single-column testing system equipped with a 1 kN load cell. 1 mL and 2.25 mL prefilled syringe (PFS) units were filled with the desired volume of test fluid and assembled with the appropriate plunger rod. Before testing, each syringe was visually inspected to confirm that it was intact and free of visible cracks and foreign matter, then secured into a universal syringe holder. The sample size was n = 3 PFS samples per analytical run. The test parameters shown in Table 1 were applied for all break loose and glide force testing. Individual measurements included break-loose force and maximum and average glide forces. Mean and standard deviation were calculated from individual-sample measurements for each viscosity sample and syringe configuration.

**Table 1.** Method Settings for Break-Loose and Glide-Force Testing and Constant-force Injection Time Testing.

| Parameter | Break-Loose and Glide-Force Test | Constant Force Injection Time Test |
| --- | --- | --- |
| Preload | 0.1 N | 0.1 N |
| Test or ramp speed | 250 mm/min | 100–1000 mm/min |
| Force hold | Not applicable | 20 N |
| End condition | 100 N upper-force limit | 100 N upper-force limit |
| Data acquisition | 100 Hz | 100 Hz |

### Injection Time at Constant Force Testing

Injection-time testing was conducted using an Instron 68SC-1 single-column testing system equipped with a 1 kN load cell. Before testing, each syringe was visually inspected to confirm that it was intact and free of visible cracks and foreign matter, then secured into a universal syringe holder. The sample size was n = 3 PFS samples per analytical run. The constant-force injection-time test parameters shown in Table 1 were applied to all injection-time measurements. For constant-force testing, an initial ramp was applied until the specified force hold of 20 N was achieved. Once the force response reached a stable plateau, the average plunger velocity was determined from the displacement-versus-time data. Injection time was calculated using the measured fluid injection length and the average velocity during the constant-force portion of the test:

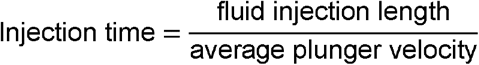

### Commercial-product laboratory assessment

One Voyxact (sibeprenlimab-szsi) prefilled syringe was assessed for the effects of product viscosity, delivered volume, and syringe geometry on injectability. BLGF was performed on the combination product using the method described above. The viscosity was then measured on the expelled drug product at room temperature and 5°C. For constant-force injection-time testing, the commercial drug product was transferred into the specified 2.25 mL test syringe to obtain an equivalent syringe and needle configuration. Results from this single-unit assessment were interpreted descriptively.

## Results

### Formulation Development

As part of the povetacicept drug product development, a high throughput, formulation screening approach was used to determine new lead formulations for further evaluation. The formulation development focused on a liquid formulation at 160-175 mg/mL for a targeted storage at 2-8°C. Variables used to address potential viscosity and stability issues included protein concentration, buffer type, formulation pH, different stabilizers and surfactants, as well as the concentrations of those constituents.

During formulation screening, two rounds of short-term stability and viscosity characterization were conducted on 20 formulations at 160 mg/mL, for a total of 40 formulations evaluated. A final, third round examined the viscosity curves of several formulations to determine if the protein concentration could be increased beyond 160 mg/mL. As a result of these three rounds, lead formulations were selected for long term stability at 160 to 175 mg/mL. As the emphasis of this paper is on the drug product viscosity and due to the proprietary nature of povetacicept; specific components, component concentrations, and other characterization data is not provided.

Table 2 shows the viscosity comparison for the first round (R1) of formulation screening, which ranged from 5.7 to 6.5 cP. A subset of four formulations with favorable stability profiles were also measured for viscosity (n=1, 20°C) after 8 weeks at 2-8°C and 30°C, with no appreciable difference between the two temperatures. The second round (R2) formulations were also evaluated for viscosity (Table 2) and ranged from 5.2 to 7.3 cP. Based on the viscosity results and associated stability over 8 weeks, two lead formulations were evaluated in Round 3 (R3), measuring viscosity over increasing protein concentrations (Figure 1 and Appendix Table A1). As is typical for protein formulations^4,5^, increases in viscosity were observed for both formulations as protein concentrations increased. At approximately 175 mg/mL, both lead formulations remained below the prespecified development target of ≤10 cP. This viscosity target was selected to ensure feasibility of the ultrafiltration/diafiltration manufacturing operation, while supporting the intended syringe configuration, injection-force objective, and rapid delivery under the bench-test conditions described below.

**Table 2.**
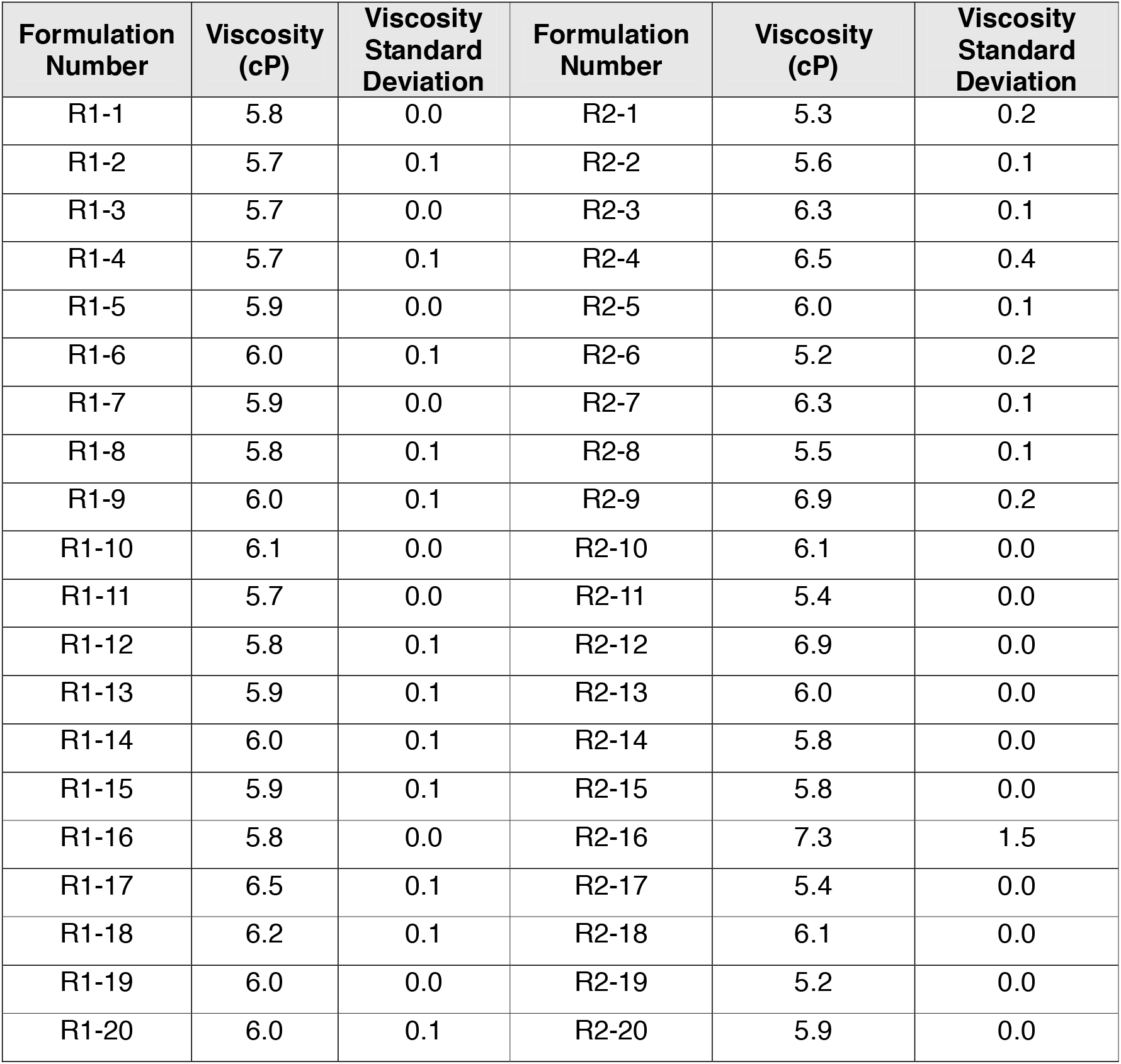
Viscosity at 20°C of Round 1 (R1) and Round 2 (R2) Formulations at 160 mg/mL.

| Formulation Number | Viscosity (cP) | Viscosity Standard Deviation | Formulation Number | Viscosity (cP) | Viscosity Standard Deviation |
| --- | --- | --- | --- | --- | --- |
| R1-1 | 5.8 | 0.0 | R2-1 | 5.3 | 0.2 |
| R1-2 | 5.7 | 0.1 | R2-2 | 5.6 | 0.1 |
| R1-3 | 5.7 | 0.0 | R2-3 | 6.3 | 0.1 |
| R1-4 | 5.7 | 0.1 | R2-4 | 6.5 | 0.4 |
| R1-5 | 5.9 | 0.0 | R2-5 | 6.0 | 0.1 |
| R1-6 | 6.0 | 0.1 | R2-6 | 5.2 | 0.2 |
| R1-7 | 5.9 | 0.0 | R2-7 | 6.3 | 0.1 |
| R1-8 | 5.8 | 0.1 | R2-8 | 5.5 | 0.1 |
| R1-9 | 6.0 | 0.1 | R2-9 | 6.9 | 0.2 |
| R1-10 | 6.1 | 0.0 | R2-10 | 6.1 | 0.0 |
| R1-11 | 5.7 | 0.0 | R2-11 | 5.4 | 0.0 |
| R1-12 | 5.8 | 0.1 | R2-12 | 6.9 | 0.0 |
| R1-13 | 5.9 | 0.1 | R2-13 | 6.0 | 0.0 |
| R1-14 | 6.0 | 0.1 | R2-14 | 5.8 | 0.0 |
| R1-15 | 5.9 | 0.1 | R2-15 | 5.8 | 0.0 |
| R1-16 | 5.8 | 0.0 | R2-16 | 7.3 | 1.5 |
| R1-17 | 6.5 | 0.1 | R2-17 | 5.4 | 0.0 |
| R1-18 | 6.2 | 0.1 | R2-18 | 6.1 | 0.0 |
| R1-19 | 6.0 | 0.0 | R2-19 | 5.2 | 0.0 |
| R1-20 | 6.0 | 0.1 | R2-20 | 5.9 | 0.0 |

**Figure 1.**
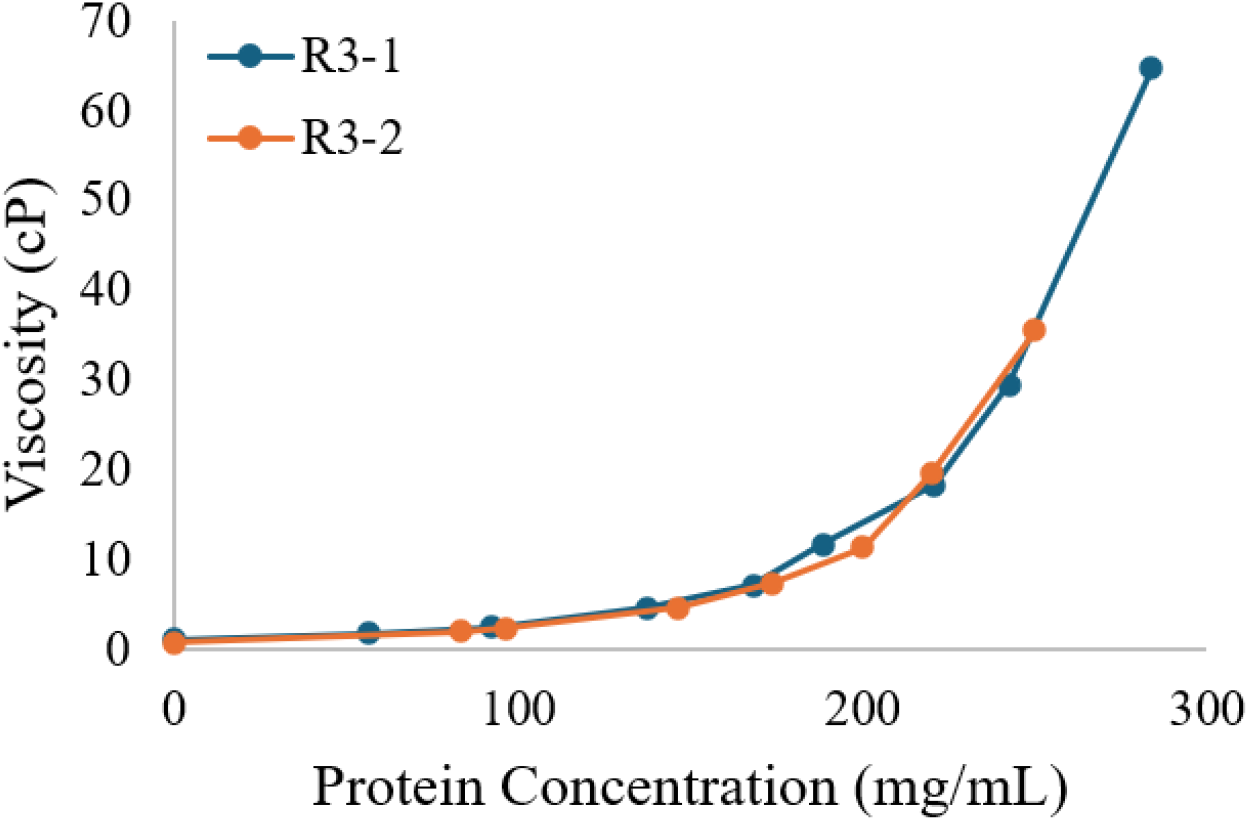
Viscosity Measurements for Lead Formulations at Varying Protein Concentrations (Round 3, 20°C, n=1)

Table 3 shows the effect of temperature on the viscosity of the final 175 mg/mL formulation selected from long-term stability and viscosity behavior. At room temperature, and even down to 15°C, the average viscosity meets the development target of ≤ 10 cP. For the treatment of IgA Nephropathy, the required dose for povetacicept is 80 mg. With these optimized drug product parameters, an injection volume of only 0.46 mL is required.

**Table 3.** Final Formulation Viscosity Measured at Various Temperatures.

| Target Temperature | Average Viscosity (cP) | Standard Deviation (cP) | Average Temperature (°C) |
| --- | --- | --- | --- |
| 5°C | 14.9 | 1.5 | 4.9 |
| 15°C | 9.8 | 0.3 | 15.0 |
| 20°C | 8.0 | 0.1 | 20.0 |
| 25°C | 6.9 | 0.1 | 25.0 |

### Pre-filled Syringe Injectability Studies

Throughout development, the povetacicept container closure evolved to a more user-friendly format for in-home use. Intentional shifts were made from vials to pre-filled syringes (PFS) and then to the intended combination product configuration using an established autoinjector platform that has been used across multiple marketed combination products. The intentional formulation choices to generate low viscosity enabled these changes.

Specifically, with the introduction of PFS, attention shifted to injectability of the product. Even with viscosity controlled to no more than 10 cP, careful attention must be paid to the dimensions of the syringe. The needle dimensions of the selected PFS must be balanced with the force required to expel the drug product. As demonstrated by R. P. Watt *et al*^1^, when the needle diameter is decreased, the maximum force required to expel the drug product increases. However, the lower gauge (larger outer diameter) needles that reduce injection force could be deleterious to the patient experience. Therefore, the commonly used 27G, thin-walled needle was selected.

The remaining variable impacting the injection force is the syringe barrel inner diameter. The 1 mL long syringe is a commonly used primary-container format for low-volume subcutaneous biologic products and was compatible with the intended 0.46 mL povetacicept dose. During dose escalation, a shift from a 1mL PFS to a 2.25mL PFS is required as the injection volume increases. Two studies were conducted to understand the injectability impact of viscosity on the two syringe sizes used for povetacicept. The first used the standard break-loose and glide force test method at a constant injection speed of 250 mm/min to measure the force required to inject. The second injected at constant forces to understand the injection time with normal patient use. As the injection force and/or duration increases, patient retention and compliance may be negatively affected.^1^ Therefore, both studies used an internal glide force target of not more than 20N informed by published literature^1^ and regulatory precedent.^13^

Figure 2 shows glide force measurements for 5, 10, 16, and 22 cP viscosity samples injected in triplicate for modeling development. The expected linear relationship between viscosity and injection force is observed, with injection force increasing with viscosity across both PFS configurations. Refer to the Appendix Table A2 for the full BLGF data summaries for 1 mL and 2.25 mL syringe configurations. When povetacicept drug product (DP) was injected under the same experimental conditions, there was good alignment with the modeling data for both syringe sizes relative to the 7 cP room temperature viscosity. The slope of the larger syringe is steeper than that of the 1 mL syringe. This, as described by Hagen-Poiseuille’s law, is due to the larger inner diameter of the syringe causing a larger volumetric flow rate through the needle at the 250 mm/min speed, which results in a larger back pressure. All experiments were conducted by expelling the test fluid into air. *In vivo* administration introduces additional tissue resistance.^14^

**Figure 2.**
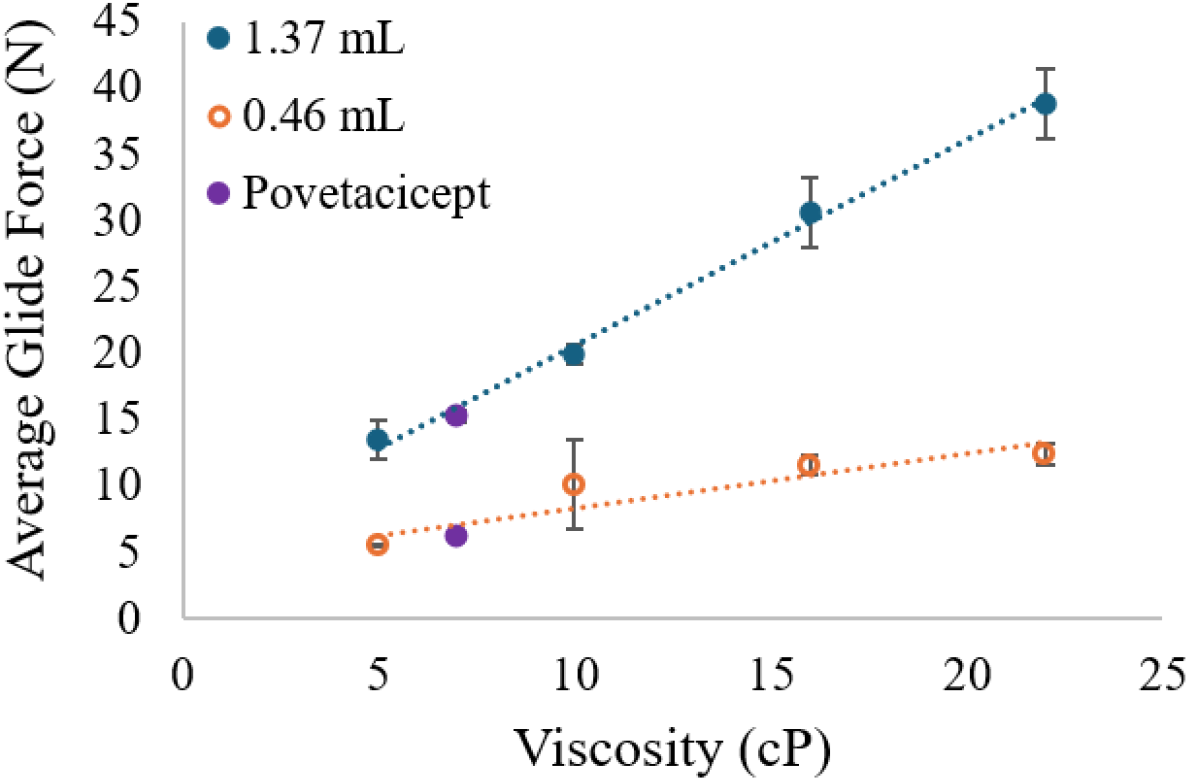
Viscosity versus Glide Force Average for 1 mL and 2.25 mL PFS.

The second series of injectability experiments focused on injection time as an output of injection force. The total work required to complete an injection includes the time the pinch force must be applied and the physical effort to maintain the force throughout the injection process.^15^ As patient-to-patient variability in injection time studies can be quite high, the injection was performed using an instrument set to maintain a constant 20N glide force. Table 4 and Table 5 show that when constant force is applied to both 1 mL and 2.25 mL PFS, there is an increase in injection time with increasing viscosity. The same dimensional factors from the Hagen-Poiseuille’s law that increase the injection force required at constant speed increases the injection time when the back-pressure is controlled by varying the volumetric flow rate. Greater flow rate reduction (injection speed) is required for 2.25 mL syringe than the 1 mL syringe. However, this relationship is not linear under these study conditions as the increase becomes more pronounced as viscosity increases. This correlates to injection forces of less-than or greater-than 20N when measured by the 250mm/min injectability study (Figure 2).

**Table 4.** Viscosity Standard Injection Time for 1 mL and 2.25 mL PFS at Constant Forces.

| Syringe Size<br>(mL) | Fill Volume<br>(mL) | Viscosity<br>(cP) | Injection Force<br>(N) | Injection Time<br>(s) |
| --- | --- | --- | --- | --- |
| 1 | 1 | 5 | 20 | 1.82<sup>a</sup> |
|  |  | 10 |  | 2.80 |
|  |  | 16 |  | 3.65 |
|  |  | 22 |  | 5.04 |
| 2.25 | 1 | 5 | 20 | 2.88 |
|  |  | 10 |  | 5.36 |
|  |  | 16 |  | 7.50 |
|  |  | 22 |  | 11.06 |

<sup>a</sup> Low viscosity samples required speeds up to 1000 mm/min. Although this is below the maximum velocity of the instrument, the short experimental duration did not allow stabilization for accurate determination of the injection velocity.

**Table 5.**
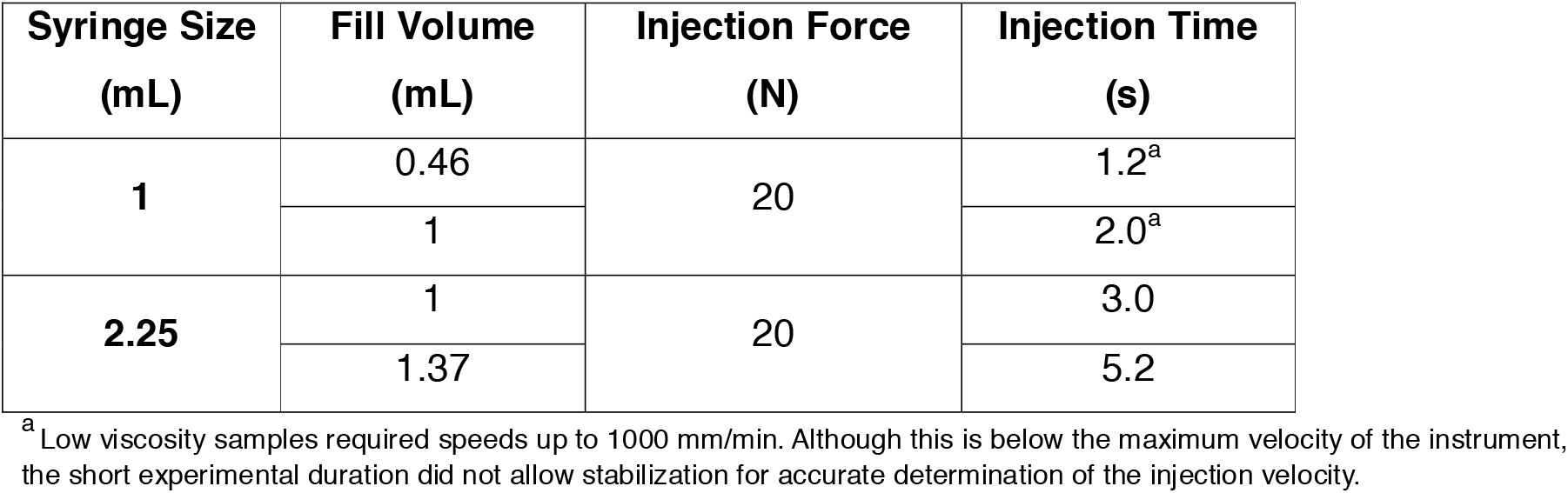
Povetacicept Injection Time for 1 mL and 2.25 mL PFS at 20N Injection Force.

| Syringe Size<br>(mL) | Fill Volume<br>(mL) | Injection Force<br>(N) | Injection Time<br>(s) |
| --- | --- | --- | --- |
| 1 | 0.46 | 20 | 1.2<sup>a</sup> |
|  | 1 |  | 2.0<sup>a</sup> |
| 2.25 | 1 | 20 | 3.0 |
|  | 1.37 |  | 5.2 |

<sup>a</sup> Low viscosity samples required speeds up to 1000 mm/min. Although this is below the maximum velocity of the instrument, the short experimental duration did not allow stabilization for accurate determination of the injection velocity.

Bringing the results of these two studies together, the ≤ 10 cP viscosity target for the povetacicept formulation provides a patient experience advantage in both injection force and injection time. Additionally, as shown in Table 5, the low volume (0.46 mL) required for the 80 mg dose enables the use of the 1 mL syringe, which provides an advantage in both injection force and time when compared to the larger syringe. However, even with the 3x dose (240 mg) in the 2.25 mL syringe, the injection force and injection time meet our internal targets of ≤ 20 N and ≤ 10 s, respectively.

A single Voyxact (sibeprenlimab-szsi) prefilled syringe was evaluated to provide descriptive external context for the bench-test relationships observed with the viscosity standards and povetacicept configurations. The Voyxact (sibeprenlimab-szsi) 2 mL dose is delivered via a pre-filled syringe with a needle safety device. The viscosity of the product was 15 cP at room temperature and 40 cP at 5°C. The average glide force was measured to be 27 N. After transfer of the drug product into the specified 2.25 mL test syringe, the calculated injection time at a constant force of 20 N was 19 seconds. These findings were consistent with the findings in this study of the relationships between viscosity, delivered volume, syringe configuration, injection force, and injection time. Because the assessment used a single commercial unit and included transfer into a test syringe for the injection-time measurement, and was conducted in air, the results should not be interpreted as a direct comparison of clinical performance, usability, or patient experience between products.

### Introduction of the Autoinjector

The patient-centered development of povetacicept did not end with a low volume injectable PFS with favorable injectability. Self-administration of the drug product by an autoinjector presented an attractive option for improved ease-of-use. Instead of a manual pinch force that must be maintained throughout the injection via PFS, a force is applied to activate the AI device and hold it in place during injection. This can lead to easier use by those with dexterity limitations. A recent qualitative survey study of adult patients with IgAN in the US revealed that 96% of those interviewed (N=24 of 25) preferred an autoinjector relative to injection via a prefilled syringe (PFS), and 80% (N=20 of 25) preferred Q4W versus Q1W dosing.^11^ Although autoinjectors ease the injection of higher viscosity drug product, higher volumes could increase the injection time and tissue back-pressure. Transition to an established autoinjector platform utilizing the same syringe characterized herein was enabled by the highly compatible low viscosity and the low volume injection. Studies of the injection force and injection time at constant forces contributed to autoinjector and drive mechanism selection. The resulting injection time for the autoinjector is measured to be 1-2 seconds in air, which is similar to the constant force injection time study results. Internal human factors studies further confirmed the patient-centric design.

## Conclusion

The povetacicept combination product and device development journey demonstrates how patient-centric development considerations begin with early formulation development through selection of the primary container and autoinjector. Intentional formulation choices, with an emphasis on low viscosity (≤10 cP) at high protein concentrations (175 mg/mL), enabled a lower volume injection (80 mg dose in a 0.46 mL volume). Low viscosity (≤10 cP) combined with the 27G ½-inch needle on a 1 mL or 2.25 mL syringe enabled low glide force (< 20 N) and faster injection time (< 10s) targets to be met. The same 1 mL syringe was incorporated into an established autoinjector platform intended for Q4W self-administration of povetacicept. The limited assessment of the commercial-product (Voyxact) provided additional context for the relationships among viscosity, dose volume, syringe configuration, injection force, and injection time.

Overall, the povetacicept development approach illustrates how dosing frequency, injection volume, formulation viscosity, delivery time, and device usability can be considered together when designing a combination product for chronic diseases such as IgAN. Taken together, these intentional choices in drug and device development may impact treatment choice, patient experience, and treatment adherence, which may potentially impact clinical outcomes.

## Appendix

**Table A1.** Viscosity Measurements for Lead Formulations at Varying Protein Concentrations (single replicate, 20°C)

| Formulation | Protein Concentration<br>(mg/mL) | Viscosity (cP) | Standard Deviation |
| --- | --- | --- | --- |
| R3-1 | 0 | 1.2 | 0 |
|  | 56.7 | 1.9 | 0.1 |
|  | 92.3 | 2.6 | 0.1 |
|  | 137.6 | 4.7 | 0.1 |
|  | 168.1 | 7.2 | 0.0 |
|  | 188.5 | 11.8 | 0.0 |
|  | 220.7 | 18.4 | 0.5 |
|  | 242.5 | 29.5 | 0.1 |
|  | 283.9 | 64.7 | 0.4 |
| R3-2 | 0 | 0.9 | 0.0 |
|  | 83.4 | 2.1 | 0.0 |
|  | 96.5 | 2.4 | 0.1 |
|  | 146.6 | 4.7 | 0.0 |
|  | 173.7 | 7.5 | 0.1 |
|  | 200.0 | 11.4 | 0.1 |
|  | 219.9 | 19.6 | 0.1 |
|  | 249.8 | 35.6 | 0.2 |

**Table A2.** Summary of Average Break-Loose and Glide Force Measurements by Fill Volume and Solution Viscosity.

| **Fill Volume (mL)** | **Viscosity (cP)** | **Mean Break-Loose Force (N)** | **Mean Maximum Glide Force (N)** | **Mean Average Glide Force (N)** |
| --- | --- | --- | --- | --- |
| **0.46** | 5 | 5.30 | 5.90 | 5.50 |
|  | 10 | 4.90 | 10.20 | 10.10 |
|  | 16 | 5.40 | 10.98 | 11.58 |
|  | 22 | 5.25 | 12.64 | 12.38 |
|  | 7 (povetacicept) | 4.60 | 8.46 | 6.24 |
| **1.37** | 5 | 5.83 | 13.50 | 13.49 |
|  | 10 | 5.88 | 19.31 | 19.96 |
|  | 16 | 6.09 | 28.86 | 30.68 |
|  | 22 | 6.09 | 37.23 | 38.80 |
|  | 7 (povetacicept) | 5.53 | 19.14 | 15.29 |

## Notes

### Competing Interest Statement

All authors received support for the present manuscript from Vertex Pharmaceuticals Incorporated.

### Funding

- Vertex Pharmaceuticals Incorporated (United States)

## References

1. Watt, R. P.; Khatri, H.; Dibble A. R. G. Injectability as a function of viscosity and dosing materials for subcutaneous administration. Int. J. Pharm. 2019, 554, 376–386.

2. Berteau, C.; Filipe-Santos, O.; Wang T.; Rojas H.E.; Granger C.; Schwarzenbach, F. Evaluation of the impact of viscosity, injection volume, and injection flow rate on subcutaneous injection tolerance. Méd. Devices: Évid. Res. 2015, 8, 473–484.

3. Jackisch, C.; Müller, V.; Maintz, C.; Hell, S.; Ataseven, B. Subcutaneous Administration of Monoclonal Antibodies in Oncology. Geburtshilfe Frauenheilkd. 2014, 74, 343–349.

4. Jezek, J.; Rides, M.; Derham, B.; Moore, J.; Cerasoli, E.; Simler, R.; Perez-Ramirez, B. Viscosity of concentrated therapeutic protein compositions. Adv. Drug Deliv. Rev. 2011, 63, 1107–1117.

5. Whitaker, N.; Xiong, J.; Pace, S. E.; Kumar, V.; Middaugh, C. R.; Joshi, S. B.; Volkin, D. B. A Formulation Development Approach to Identify and Select Stable Ultra-High-Concentration Monoclonal Antibody Formulations With Reduced Viscosities. J. Pharm. Sci. 2017, 106 (11), 3230–3241.

6. Jayaprakash, V.; Costalonga, M.; Dhulipala, S.; Varanasi, K. K. Enhancing the Injectability of High Concentration Drug Formulations Using Core Annular Flows. Adv. Healthc. Mater. 2020, 9, e2001022.

7. Zhao, Y.; Ma, S. Observations on the Prevalence, Characteristics, and Effects of Self-Treatment. Front. Public Heal. 2016, 4, 69.

8. Freundlich, B.; Kivitz, A.; Jaffe, J.S. Nearly Pain-Free Self-administration of Subcutaneous Methotrexate with an Autoinjector. JCR: J. Clin. Rheumatol. 2014, 20, 256–260.

9. Dostal, P.; Taubel, J.; Lorch, U.; Aggarwal, V.; York, T. The Reliability of Auto-Injectors in Clinical Use: A Systematic Review. Cureus. 2023, 15, e41601.

10. Bowen, M.; Adam, Y.; Chang, D. Advancements and Challenges in Large Molecule Drug Delivery and Devices for Patient-Centric Care. AAPS Open. 2026, 12, 50.

11. Iff, J.; Mellor, R.; Nair, R.; Maski, M.; Lin, W.; Liu, J.; Wu, C-H,; Poulos, C. Patient Perceptions of Injectable Therapies Among Adults with IGAN: A Qualitative Study. Presented at the American Society of Nephrology Kidney Week, Denver, CO, USA, October 21–25, 2026.

12. Madan, A.; Yalavarthy, R.; Kim, D. K.; Moon, J-Y.; Park, I.; Mandayam, S.; Cortazar, F.; Kim, S. G.; Enstrom, A.; Thomas, H. et al. Povetacicept for IgA Nephropathy and Primary Membranous Nephropathy. Kidney Int. Rep. 2026, 11, 106–116.

13. FDA Center for Drug Evaluation and Research. Application Number: 761111Orig1s000; Reference ID: 4631680; 2020. https://www.accessdata.fda.gov/drugsatfda_docs/nda/2020/761111Orig1s000OtherR.pdf

14. Allmendinger, A.; Mueller, R.; Schwarb, E.; Chipperfield, M.; Huwyler, J.; Mahler, H-C.; Fischer, S. Measuring Tissue Back-Pressure – In Vivo Injection Forces During Subcutaneous Injection. Pharm. Res. 2015, 32, 2229–2240.

15. Zhang, Q.; Fassihi, M. A.; Fassihi, R. Delivery Considerations of Highly Viscous Polymeric Fluids Mimicking Concentrated Biopharmaceuticals: Assessment of Injectability via Measurement of Total Work Done “WT”. AAPS PharmSciTech. 2018, 19, 1520–1528.

